# Trophic interactions in microbiomes influence plant host population size and ecosystem function

**DOI:** 10.1101/2023.03.06.531362

**Authors:** Jiaqi Tan, Na Wei, Martin M. Turcotte

## Abstract

Plant microbiomes that comprise diverse microorganisms, including prokaryotes, eukaryotes, and viruses are the key determinant of plant population dynamics and ecosystem function. Despite their importance, little is known about how interactions, especially trophic interactions, between microbes from different domains modify the importance of microbiomes for plant hosts and ecosystems. Using the common duckweed *Lemna minor*, we experimentally examined the effects of predation (by bacterivorous protozoa) and parasitism (by bacteriophage) within microbiomes on plant population size and ecosystem phosphorus removal. Our results revealed that predation increased plant population size and phosphorus removal whereas parasitism showed the opposite pattern. The structural equation modeling further pointed out that predation and parasitism affected plant population size and ecosystem function via distinct mechanisms that were both mediated by microbiomes. Our results highlight the importance of understanding microbial trophic interactions for predicting the outcomes and ecosystem impacts of plant-microbiome symbiosis.

## Introduction

Plants are key to many ecological communities, driving critical ecosystem functions, such as primary production and nutrient cycling [1–3]. The central role of plants in mediating ecosystem functioning is increasingly recognized to be influenced by associated microbiomes (i.e., plant microbiomes) [4–6]. Indeed, diverse microorganisms within plant microbiomes can potentially interact with each other via different types of interactions and have significant impacts on plants and ecosystem functioning [7–9]. While it is well established that microbiomes can be directly involved in ecosystem functions or indirectly by affecting plant hosts’ physiology, growth, and reproduction, the extent to which species interactions within plant microbiomes (microbe-microbe interactions) affect plant populations and ecosystem functions remains unclear [4, 10, 11]. As microbe-microbe interactions have often been studied in pairs or synthetic communities in vitro [10, 12, 13], they often do not fully capture the complexity of natural microbiomes that comprise producers, consumers, decomposers, and more. Ignoring the emergent impacts of common species interactions within microbiomes may limit our understanding of how microbiomes influence their macroscopic hosts and ecosystem functions.

Trophic interactions are ubiquitous in nature [14–17]. For example, macroscopic predators, such as soil nematodes and mosquito larvae, have been shown to affect the diversity and composition of soil and plant microbiomes [18–21]. Although knowledge of interactions between metazoan predators and microbial prey suggests the relevance of trophic interactions in influencing microbiomes, little is known about the role of trophic interactions between microbes within plant microbiomes in mediating microbiome structure and function on plant population dynamics and ecosystem function. In plant microbiomes, prokaryotes, viruses, and eukaryotes form complex networks [7, 22] via horizontal resource-mediated interactions (e.g., competition and mutualism) and via trophic interactions that vertically connect microbial hosts with parasites and predators with prey (e.g., parasitism and predation) [8]. Among the interacting microbes, bacteriophages [23] and protozoa [24] are two major types of bacterivorous consumers (parasites and predators, respectively). These microbial consumers may directly influence plants and ecosystem functions themselves, or indirectly by affecting bacteria in the microbiomes. For instance, bacteriophages and protozoa themselves may influence plant populations and ecosystem functions if they have the ability to recycle limiting nutrients and secrete growth-stimulating substance to the environment [25]. Bacteriophages and protozoa through trophic interactions with bacteria may also have the potential to alter the properties of microbiomes (e.g., microbial abundance, richness, and composition), translating into an impact on host plant populations and ecosystem functions [7].

While both being bacterivorous consumers, bacteriophages and protozoa may exhibit distinct impacts on microbiomes as microbial parasites vs. predators [24, 26–28]. For instance, these two types of consumers may show considerable variation in diet breath. Bacteriophages are highly specialized and can specifically target certain groups of microbes [23, 29], such as plant growth-promoting or growth-suppressing bacteria. As such, they may strongly affect the richness and composition of microbiomes and the directionality (positive or negative) of microbiome effects on plant populations. By contrast, protozoa are often generalists and can capture a wide variety of bacteria through endocytosis, leading to a reduction in the overall microbial abundance and thereby potentially influencing the functions of microbiomes on plants [24]. Despite the prevalence and significance of microbial bacterivorous consumers in plant microbiomes, little is known about how they impact microbiomes and how the impacts translate into changes in plant populations and ecosystem functions. Specifically, while bacteriophages and protozoa have been studied separately [23, 26, 28, 29], their influences on plant hosts and ecosystem functions have not been directly compared.

Here we examined how trophic interactions within plant microbiomes influenced plant populations and ecosystem functions using the common duckweed, *Lemna minor*, and their natural microbiomes. By experimentally manipulating parasitism (by bacteriophages) and predation (by protozoa) in the bacterial microbiomes of *L. minor*, we examined the effects of trophic interactions on bacterial microbiomes and plant population size and phosphorus removal, a key ecosystem function performed by microbes and plants in freshwater ecosystems.

## Materials and Methods

### Study system and sampling

We used *Lemna minor* as the plant host for our experiment. Two *L. minor* genotypes that were genetically differentiated by two polymorphic microsatellite loci [30, 31] were collected from lakes in Boyce Mayview Park (40.33566°N, 80.11269°W) and Deer Lakes State Park (40.62245°N, 79.82246°W) in Pennsylvania, USA. To create axenic *L. minor* lines, we first rinsed *L. minor* in 4.5 ml sterile water in glass test tubes (16 × 150 mm) and then transferred into 8 ml 0.025 M phosphate buffer saline (PBS) and sonicated for 5 min. We then bleached *L. minor* with 1% sodium hypochlorite for 20 s before transferring into sterile media (8 ml 1:4 Schenk and Hildebrandt (SH) basal salt medium; Sigma-Aldrich, Inc., St. Louis, Missouri, USA) for propagation under 24 h light. The axenic cultures of *L. minor* were verified using PCRs with bacterial 16S and fungal internal transcribed spacer to confirm that the absence of epiphytic and endophytic microbes [10].

In this study, we used bacterial microbiomes of duckweeds that were naïve to the two focal *L. minor* genotypes to avoid pre-existing local adaptation of microbiomes to host plants that might cause genotypic differences in fitness. Bacterial communities and bacteriophages were extracted from the natural microbiomes of *L. minor* collected from the City Park Lake (30.42900°N, 91.16747°W; microbiome source 1) and College Lake (30.40714°N, 91.16888°W; microbiome source 2) in Louisiana, USA. To do so, we collected 50 clusters of *L. minor* (*c*. 150 individuals) at each location and vortexed them gently in 4.5 ml sterile water for 10 s to remove the microbial residents from lake water. We then transferred the clusters to 8 ml PBS and sonicated at 40 kHz for 5 min to release duckweed microbiomes into PBS.

We removed the eukaryotic microbes from the microbiomes and isolated bacteria and bacteriophages separately. To obtain the bacteria, we first filtered 6 ml of the microbiome extracts using 1.2 μm PTFE sterile syringe filters (Acrodisc, Pall Laboratory, New York, USA) to remove eukaryotes, and confirmed the removal of eukaryotes by screening 0.1 ml of the filtered microbiome extracts using optical microscopy. We then pelleted bacterial cells by centrifuging at 3,000 rpm for 15 min. The supernatant containing most bacteriophages was discarded. Given the possibility that a small number of bacteriophages might remain in the bacterial pellets, we further purified the bacterial isolates by suspending the bacterial pellets in 1 ml PBS, shaking at 500 rpm for 30 min, and centrifuged at 3,000 rpm for 15 min to concentrate the bacterial pellets in 0.05 ml PBS. These steps were repeated six times, resulting in ~ 6.4 × 10^7^ times dilution of bacteriophages in bacterial cell pellets. The final bacterial pellets were suspended in 1 ml 15% glycerol solutions and preserved at −80°C.

To isolate the bacteriophages, we filtered 2 ml of the original microbiome extracts using 1.2 μm (Acrodisc, Pall Biotech, Westborough, Massachuttes, USA) and then 0.2 μm (Whatman Uniflo, Cytiava, Massachuttes, USA) PTFE sterile syringe filters. The presence of phage particles was confirmed by screening 10 μl of the filtered extracts using epifluorescence microscopy [32, 33]. Phage isolates were preserved at −80 °C.

We found several ciliated protozoa (bacterivorous filter feeders) in the *L. minor* microbiomes, and chose a common ciliate, *Tetrahymena pyriformis*, as a representative of the ciliated protozoa communities since protozoa are generalist predators often with similarly broad feeding preferences [34]. We acquired axenic *T. pyriformis* from the Carolina Biological Supply (Burlington, North Carolina, USA), and maintained it in 6 ml 10 g/L peptone water at 24 °C prior to the experiment.

### Trophic interaction experiment

We used a three-way factorial design that manipulated host plant genotypes (A and B), microbiome sources (1 and 2), and trophic interactions (control, parasitism by bacteriophages, and predation by protozoa). We replicated the experiment three times with 36 total microcosms. These microcosms were in 8 ml autoclaved 1:4 SH salt medium in loosely capped glass test tubes (16 × 150 mm).

For the experiment, we first introduced one cluster (with three to four individuals) of axenic *L. minor* into each microcosm. We then inoculated the bacterial microbiomes by thawing the frozen bacterial isolates, diluting in 8 ml 1:4 SH salt medium, and transferring 0.1 ml of ~10^5^ cells/ml into the designated microcosms. In the parasitism treatment, we thawed the frozen virus isolates and transferred 10 μl of the isolates to the designed microcosms. The virus isolates from microbiome sources 1 and 2 contained 520 and 240 virus particles (in 10 μl), respectively, and were added to microcosms that contained the bacteria from the same source. In the predation treatment, we diluted the axenic culture of *T. pyriformis* in 1:4 SH salt medium and inoculated ~100 individuals (10 μl) into each of the designated microcosms. The control treatment received 10 μl autoclaved 1:4 SH salt medium instead. The microcosm experiment was carried out in a Percival light incubator (I-36LL; Percival Scientific, Perry, Iowa, USA) under the 12 h photocycle and 24 °C /16 °C day/night temperatures. The photosynthetic photon flux level was maintained at 75 μmol m^−2^ s^−1^ during the day. Microcosms were randomized once a week. As *L. minor* populations reached equilibrium in four weeks, the experiment was terminated on day 35.

To further examine whether parasitism and predation treatments directly affect *L. minor* population size and ecosystem function in the absence of bacterial microbiomes, we set up a follow-up experiment where we manipulated host genotypes (A and B) and trophic interaction types (control, parasitism, and predation) without bacterial microbiomes. This follow-up experiment was also replicated three times and followed the settings as described above.

### Plant population size and phosphorus concentration in the microcosms

We measured plant population size by counting the number of *L. minor* individuals in each microcosm at the end of the experiment. For phosphorus concentration, we measured total phosphorus concentration remained in the medium, following Wetzel and Likens [35]. As each microcosm started with the same amount of phosphorus at the beginning of the experiment, a lower remaining phosphorus concentration indicated a higher phosphorus removal rate by organisms in the microcosms. Briefly, we gently vortexed the microcosms, collected 1 ml from each microcosm, and removed particles (such as plant tissues and bacteria and protist cells) by filtration through 0.2 μm PTFE filters (Whatman Uniflo). The phosphorus in the 1 ml sample was first oxidized by 0.16 ml 5% persulfate solution and then reacted with 0.1 ml composite reagent of 0.6% ammonium molybdate, 7.5% sulfuric acid, 1.2% ascorbic acid, and 0.014% potassium antimonyl-tartrate, prior to reading under OD885 against the standard solutions of potassium dihydrogen phosphate (0-2 mg/L) using a microplate reader (EPOCH2, Biotek, Santa Clara, California, USA).

### Bacterial microbiomes of the microcosms

We measured the bacterial abundance, richness, and community composition in each microcosm at the end of the experiment. To measure bacterial abundance, we sonicated the microcosms for 5 min to release duckweed microbiomes to the liquid. Due to the presence of other particles (e.g., plant tissues and protist cells) in the microcosms, we stained the bacterial cells in a 0.15 ml sample with 5% crystal violet. The cells were then pelleted and rinsed twice with 95% ethanol and screened under OD570. We approximated bacterial cell concentration as ~8 × 10^8^ cells/ml per one OD570 unit.

To measure bacterial richness and community composition, we conducted microbiome sequencing. To do so, we pelleted the bacterial cells by centrifuging (3,000 rpm for 30 min) a 4.5 ml sample of each microcosm and extracted bacterial DNA with a Quick-DNA Fecal/Soil Microbe Miniprep Kits (Zymo Research, Irvine, California, USA). Two negative controls that contained the SH salt medium alone were included in the process of DNA extraction. The 36 samples and two negative controls were sent to Novogene Corporation (Sacramento, California, USA) for library preparation (16S rRNA V5–V6 region) and sequencing on an Illumina MiSeq paired-end 250 bp. While the two negative controls failed in sequencing (too few reads), we obtained 81,913 raw reads on average per sample (range = 51,411–98,957).

These paired-end (PE) raw reads were used for detecting bacterial amplicon sequence variants (ASVs) using the package DADA2 v1.14.0 [36] in R v3.6.2 [37]. Following previous pipelines [38, 39], the PE reads were trimmed and quality filtered [truncLen = c(220, 220), maxN = 0, truncQ = 2, maxEE = c(2,2)], prior to specific variant identification that took into account sequence errors. The PE reads were then end joined (minOverlap = 20, maxMismatch = 4) for ASV detection and chimera removal. ASVs were assigned with taxonomic identification based on the SILVA reference database (132 release NR 99) implemented in DADA2. A rooted phylogenetic tree of the ASVs was built using QIIME 2 v2019.10 [40]. These ASVs were further filtered using the package phyloseq [41]. First, we removed non-focal ASVs (Archaea, chloroplasts, and mitochondria). Second, we conducted rarefaction analysis using the package iNEXT [42] to confirm that the sequencing effort was sufficient to capture the bacterial richness (Fig. S1). We further normalized per-sample reads by rarefying to the lowest number of the filtered reads (28,343) across samples. Lastly, we removed low-frequency ASVs (<0.001% of total observations). The final bacterial community matrix consisted of 497 ASVs across the 36 samples.

### Statistical analyses of plant population size and phosphorus concentration

To evaluate how trophic interactions, microbiome sources, and host plant genotypes influence plant population size and total phosphorus concentration, we conducted general linear models (LMs) in R. The predicators of the LMs included trophic interaction (control, predation, and parasitism), microbiome source (1 and 2), and host genotype (A and B), as well as their two-way and three-way interactions. The response variables were natural log transformed to improve normality. The data from the main and follow-up experiment (in the absence of bacterial microbiomes) were analyzed separately. Statistical significance (type III sums of squares) and least-squares means (LS-means) of predictors were assessed using the packages emmeans [43] and phia [44].

### Statistical analyses of bacterial abundance, richness, and community composition

To evaluate how trophic interactions, microbiome sources, and host plant genotypes influence bacterial abundance (OD570) and richness (ASVs), the predictors of the LMs included trophic interaction (control, predation, and parasitism), microbiome source (1 and 2), and host genotype (A and B), as well as their two-way and three-way interactions. The response variable (bacterial abundance or richness) of each LM was power transformed if necessary to improve normality, with the optimal power parameter determined using the Box–Cox method in the package car [45] (power parameter = 1, no transformation, for both bacterial abundance and richness here).

Bacterial community composition was assessed based on Bray–Curtis dissimilarity using constrained principal coordinates analysis (cPCoA) in the package vegan [46]. To assess the significance of the main effects, the predicators of cPCoA included trophic interaction, microbiome source, and host genotype. To assess the significance of two-way interactions, the predicators of cPCoA included both the main effects and their two-way interactions. Likewise, to assess the significance of the three-way interaction, the predicators of cPCoA included the main effects, two-way interactions, and the three-way interaction following our previous work [47].

To further pinpoint the bacterial ASVs within the microbiomes that were affected by trophic interactions as shown in the LM and cPCoA as well as the structural equation modeling below, we used the packages randomForest [48] and edgeR [49]. Specifically, as the predation treatment reduced bacterial abundance relative to the control (Figs. 2 and 3), we used random forest classification to identify the ASVs that differed between the predation and control treatments with lower absolute abundances in the predation treatment. To do this, the random forest classification used the absolute abundances of the ASVs, which were the product of the relative abundances of ASVs within a microbiome and the total bacterial cells of a microcosm (~8 × 10^8^ cells/mL × OD570 × 8 mL microcosm). The random forest classification models were run for the full data set with 1000 trees, and the number of randomly selected variables (i.e., individual ASVs) at each split of a decision tree was optimized using 10-fold cross validation in the package caret [50]. Model performance was assessed using out-of-bag (OOB) error. The set of ASVs that differed between the predation treatment and the control were selected using backward variable elimination with the package varSelRF [51]. Different from bacterial abundance, bacterial richness and composition (cPCoA 2) were influenced by the parasitism treatment (Figs. 2 and 3). As bacterial richness was based on the rarefied bacterial community matrix (i.e. relative abundances), we used edgeR that was designed for modeling relative abundances using generalized linear models (GLMs) with negative binomial errors [49], to identify the ASVs that were significantly reduced in the parasitism treatment relative to the control. Our design matrix followed host genotype + microbiome source + trophic interaction to account for the potential influence of confounding factors. The model was fitted using glmQLFit() and specific contrasts were made by glmQLFTest(). The ASVs that were significantly reduced in the parasitism treatment relative to the control were identified with a false discovery rate (FDR) < 0.05. For bacterial community composition (cPCoA 2), we focused on the ASVs with large influence (scores ≥ 0.01 or ≤ −0.01) on the cPCoA 2.

### Structural equation modeling (SEM)

To evaluate how trophic interactions influence plant population size and ecosystem functioning via bacterial microbiomes, we conducted SEM to link trophic interactions to bacterial abundance (OD570), richness (ASVs), and community composition (cPCoA 2, which was strongly influenced by trophic interactions; Fig. 2c) to plant population size and total phosphorus concentration. The SEM did not include microbiome source (1 and 2) as an exogenous variable, because microbiome source showed weak effects on bacterial abundance, richness, and bacterial cPCoA 2, although it influenced bacterial cPCoA 1 (Fig. 2). Likewise, the SEM did not include host genotype (A and B) as an exogenous variable, due to its weak effects on bacterial microbiomes (abundance, richness, and composition) and plant population size, especially compared to trophic interactions (Figs. 1 and 2). For trophic interactions, we coded the control treatment as the reference level so that the effects of all other treatments (predation and parasitism) were relative to the control. Plant population size and total phosphorus concentration were natural log transformed as described in the LMs above. The full model of SEM (Fig. S2) was fitted using maximum likelihood with robust Huber-White standard errors using the package lavaan [52]. To reduce model complexity, the SEM was re-fitted with only notable paths (*P* < 0.10) present. Model fit was confirmed (comparative fit index, CFI > 0.9; root mean squared error of approximation, RMSEA, the lower bound of 90% confidence interval < 0.05; standardized root mean squared residual, SRMR < 0.1). To account for the potential influence of limited sample size, we further re-fitted the SEM using bootstrapping in lavaan. The two estimators (maximum likelihood and bootstrapping) resulted in consistent results.

**Fig. 1.**
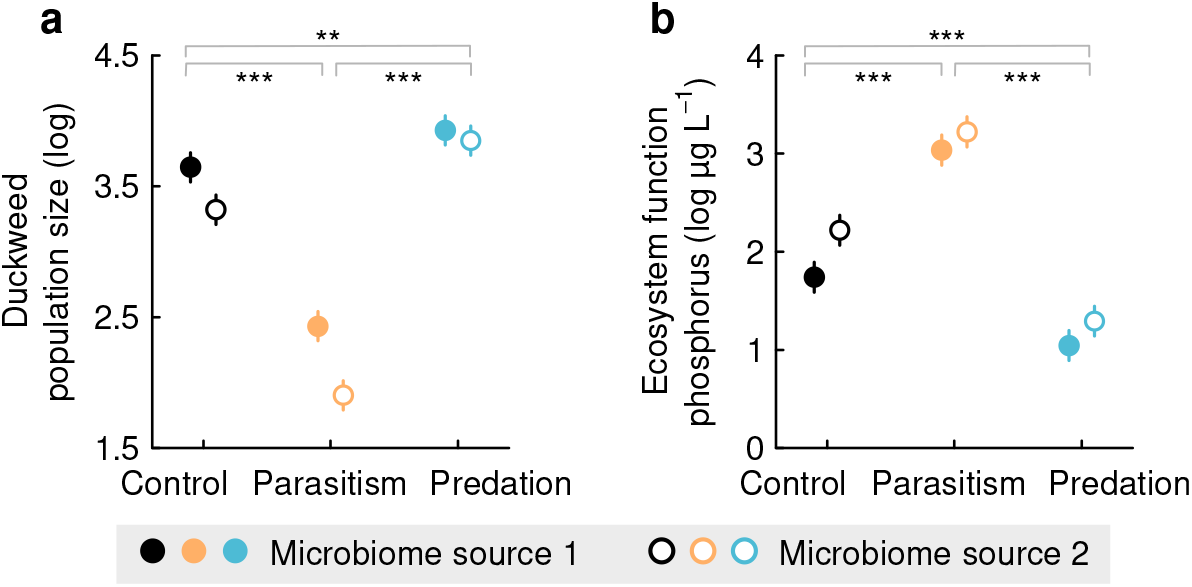
Plant population size and ecosystem function differ strongly among trophic interaction treatments. The least-squares means (LS means) of natural log transformed (**a**) duckweed population size and (**b**) ecosystem phosphorus concentration are plotted with error bars (SE) based on general linear models (predictors: interaction treatment, microbiome source, genotype, and their two-way interactions). Significant LS mean contrasts between treatments are denoted: \*\*\**P* < 0.001; \*\**P* < 0.01; \**P* < 0.05. Duckweed population size increased and phosphorus concentration decreased in the parasitism treatment relative to the control. The opposite pattern was observed in the predation treatment relative to the control. See statistical details in Tables S1 and S2.

**Fig. 2.**
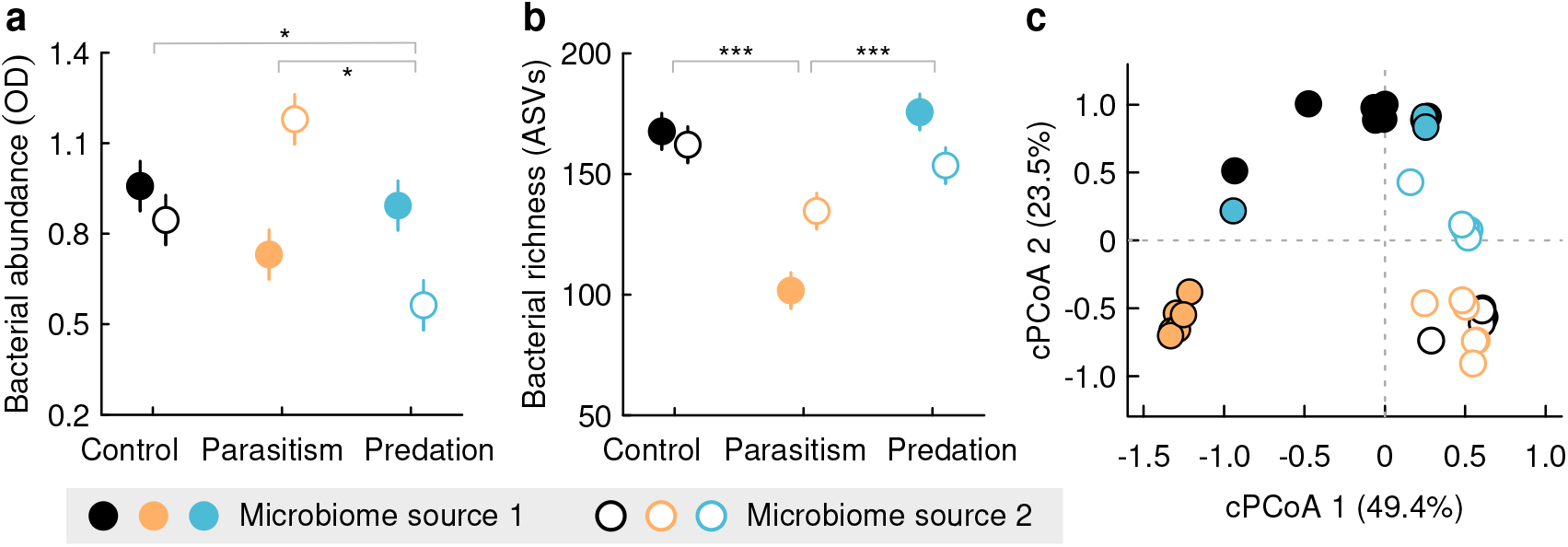
Bacterial microbiomes differ strongly among trophic interaction treatments. Bacterial microbiomes were characterized using (**a**) optical density (OD) at 570 nm to indicate bacterial abundance, (**b**) amplicon sequence variants (ASVs) to represent bacterial richness, and (**c**) constrained principal coordinates analysis (cPCoA) to characterize community composition. The least-squares means (LS means) of bacterial OD and ASVs are plotted with error bars (SE) based on general linear models (predictors: interaction treatment, microbiome source, genotype, and their two-way interactions). Significant LS mean contrasts between treatments are denoted: \*\*\**P* < 0.001; \*\**P* < 0.01; \**P* < 0.05. The cPCoA revealed a strong effect of microbiome source (*F* = 24.5, *P* = 0.001) on composition along the first axis (cPCoA 1), and a strong effect of interaction treatment (*F* = 10.6, *P* = 0.001) along the second axis (cPCoA 2). See statistical details in Tables S3, S5, and S7.

## Results

### Contrasting effects of parasitism and predation on plant population size and phosphorus concentration

Duckweed population size was significantly lowered in the parasitism treatment but increased in the predation treatment relative to the control (Fig. 1a and Table S1; LM: trophic interaction, *F* = 129.6, *P* < 0.001; planned contrasts within the LM: control vs. parasitism, *F* = 138.7, *P* < 0.001, control vs. predation, *F* = 13.1, *P* = 0.001). The opposite pattern was observed for the concentration of remaining phosphorus: higher in the parasitism treatment but lower in the predation treatment relative to the control (Fig. 1b and Table S2; trophic interaction: *F* = 82.9, *P* < 0.001; control vs. parasitism, *F* = 56.4, *P* < 0.001, control vs. predation, *F* = 28.1, *P* < 0.001). In the absence of the bacterial microbiomes, protists and phages did not influence duckweed population size (*F* = 0.20, *P* = 0.66) and phosphorus concentration (*F* = 0.11, *P* = 0.74), indicating that the effects of parasitism and predation on duckweed populations and phosphorus concentrations were mediated via the microbiomes.

### The mechanisms by which parasitism and predation influence plant population size and phosphorus concentration

To examine how trophic interactions within the microbiomes influenced duckweed population size and phosphorus concentration, we sequenced the microbiomes using 16S rRNA sequencing. We found that trophic interactions had a significant impact on bacterial abundance, richness, and community composition (Fig. 2), with parasitism and predation affecting different aspects of the microbiomes. Specifically, in contrast to parasitism, predation drove a decline in bacterial abundance (LM, *F* = 4.45, *P* = 0.045; Fig. 2a and Table S3), especially in Burkholderiaceae as revealed by the random forest classification analysis (Table S4). Parasitism instead drove a decline in bacterial richness (*F* = 39.4, *P* < 0.001; Fig. 2b and Table S5), with 25 bacterial ASVs across multiple families (e.g., Beijerinckiaceae, Burkholderiaceae, Microbacteriaceae, Mycobacteriaceae, Rhizobiaceae, Sphingomonadaceae, and Xanthobacteraceae) significantly depleted in the parasitism treatment relative to the control (Table S6). In addition, parasitism drove the separation of the community composition from the control and predation treatment along the second axis of the constrained principal coordinates analysis (cPCoA 2, trophic interaction, 27% of total variation, *F* = 10.6, *P* = 0.001; Fig. 2c and Table S7), especially in Beijerinckiaceae, Burkholderiaceae, Enterobacteriaceae, Microbacteriaceae, Mycobacteriaceae, Rhizobiaceae, Sphingomonadaceae and Xanthobacteraceae (Table S8).

We further conducted SEM to examine how such impacts of trophic interactions on different properties of microbiomes influenced duckweed population size and phosphorus concentration (Fig. 3). We found that the parasitism treatment affected duckweed population size by affecting bacterial richness and community composition (Fig. 3 and Table S9). Specifically, as shown in the LM (Fig. 2b), parasitism reduced bacterial richness (*r* = −0.71, *P* < 0.001; Fig. 3 and Table S9), and such reduction in bacterial richness had a negative effect on duckweed population size (*r* = −0.18, *P* = 0.033). As a result, parasitism showed a positive effect on plant population size by reducing bacterial richness (*r* = −0.71 × −0.18 = 0.13). By contrast, parasitism showed a negative effect on plant population size by altering bacterial community composition (*r* = −0.65 × 0.27 = −0.18). In addition to bacterial richness and community composition, parasitism treatment showed a direct negative effect on duckweed population size (*r* = −0.76, *P* < 0.001). We confirmed that this direct negative effect was not caused by phages per se (see Results above) but by other unmeasured properties of the microbiomes.

**Fig. 3.**
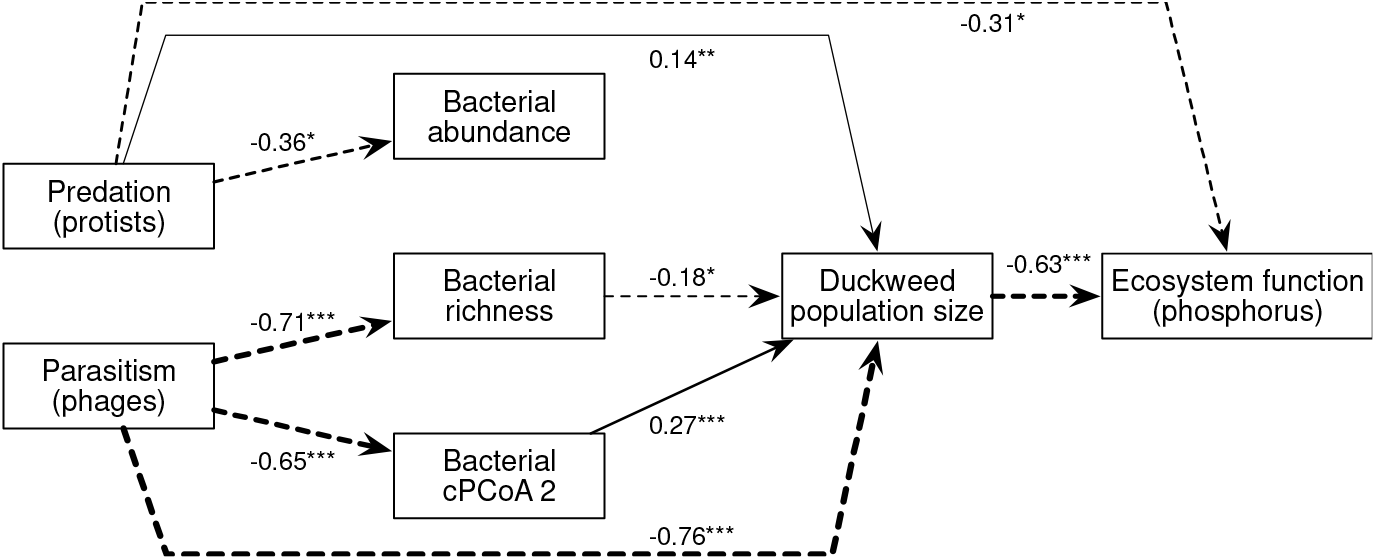
Structural equation model (SEM) of trophic interactions in microbiomes explaining duckweed population size and ecosystem function. Arrows indicate significant positive (solid) and negative (dashed) relationships (for the original full model of SEM, see Fig. S2). The SEM was estimated with both Huber–White robust standard errors (SEs, Table S9) and bootstrapping (shown here) and yielded consistent results. Numbers adjacent to arrows indicate standardized path coefficients. Trophic interactions were coded using the control treatment as the reference level in the SEM. Bacterial cPCoA 2 represented the second axis of the constrained principal coordinates analysis of bacterial community composition (Fig. 2c). Significance levels: \**P* < 0.05; \*\**P* < 0.01; \*\*\**P* < 0.001. See statistical details in Table S9.

The predation treatment affected plant population size mostly through different mechanisms than the parasitism treatment (Fig. 3). While predation reduced bacterial abundance (*r* = −0.36, *P* = 0.025; Fig. 3) as also revealed by the LM (Fig. 2a), such reduction in bacterial abundance did not significantly affect plant population size (*r* = −0.12, *P* = 0.106). Instead, predation treatment showed a direct positive effect on plant population size (*r* = 0.14, *P* = 0.006). We also confirmed that this direct positive effect was not caused by protists per se (see Results above) but by other unmeasured properties of the microbiomes.

In addition to plant population size, parasitism and predation affected phosphorus concentration via different mechanisms (Fig. 3). The parasitism treatment had a net positive effect on phosphorus concentration through reducing plant population size caused by changes in the microbiomes (*r* = (−0.76 + −0.65 × 0.27 + −0.71 × −0.18) × −0.63 = 0.51). By contrast, predation treatment had a net negative effect on phosphorus concentration (*r* = 0.14 × −0.63 + −0.31 = −0.40) through affecting plant population size (*r* = −0.63 × 0.14 = −0.09) and through microbiomes directly (*r* = −0.31, *P* = 0.015).

## Discussion

Trophic interactions shape ecological populations and communities and drive ecosystem functions [53–56]. By focusing on two important types of trophic interactions in plant microbiomes—predation (by protozoa) and parasitism (by bacteriophages)—our results revealed that predation reduced bacterial abundance, whereas parasitism reduced bacterial richness and shifted community structure. As a result of these distinct effects on microbiomes, predation and parasitism led to opposing outcomes in mediating plant population size and phosphorus removal. Specifically, predation showed a positive effect on plant population size and phosphorus removal whereas parasitism showed an opposite pattern. The SEM also pointed out the role of other mechanisms by which predation and parasitism altered microbiomes beyond abundance, richness, and composition to influence plant population size and phosphorus removal.

Our results showed that predation and parasitism influenced different properties of microbiomes. Predation by generalist protists reduced bacterial abundance, especially the most abundant ones (e.g. Burkholderiaceae; Table S4), consistent with a density-dependent foraging behavior often exhibited by generalist predators [57]. Although rarer bacteria could increase in abundance (Table S4) likely by colonizing the vacated niches of the abundant bacteria, this was yet insufficient to compensate for the loss of overall bacterial abundance, resulting in a net negative effect of predation on bacterial abundance. In contrast to predation, parasitism by specialist bacteriophages had little impact on bacterial abundance but reduced bacterial richness. Several bacteria were significantly depleted in the parasitism treatment (Table S6), likely due to the presence of different bacteriophages that target each own bacterial species in our phage extracts from the natural duckweed microbiomes. As many of these depleted bacterial species were not among the most abundant ones in the microbiomes, parasitism did not affect overall bacterial abundance. Such distinct effects of the two major consumers (microbial predators and parasites) on microbiomes contributed to our understanding of the principles that govern microbiome assembly and the roles of different types of trophic interactions.

As a result of the distinct effects on microbiomes, not only did predation and parasitism show opposite effects on plant population size but also acted via different mechanisms. For instance, predation reduced bacterial abundance, but such reduction of bacterial abundance did not translate into a significant impact on duckweed population size. Instead, predation increased duckweed population size by influencing aspects of the microbiomes (the direct link, *r* = 0.14) other than abundance, richness, and composition. It is possible that predation could induce physiological or evolutionary changes of bacteria to influence duckweed population size. For instance, predation could increase the overall activity level of the microbiomes to affect plant population size by increasing the proportion of active relative to dormant bacteria. This is because filter-feeding protists can consume dormant bacteria (different from parasitic phages that preferentially target active bacteria) [58], and the vacated niches could be occupied by active bacteria that recover more quickly than dormant bacteria from predation. Moreover, predation might drive the rapid evolution of consumer-resisting traits in bacteria, for example, forming biofilm against predation [59] that could also alter the functions of microbiomes and change plant growth [60]. In contrast to predation, parasitism showed a net negative effect on duckweed population size by influencing bacterial richness and community composition, as well as other aspects of the microbiomes. Interestingly, the reduction in bacterial richness by parasitism translated to a positive effect on duckweed populations size. The depleted species by parasitism were often rarer (Table S6) and possibly play a less important role in benefiting host plants compared to more abundant ones that could colonize the vacated niches of the deleted rarer species and benefit host plants. Moreover, the shifted community composition by parasitism led to a negative effect on duckweed population size. This was likely because parasitism changed the composition of many bacteria that belong to the bacterial groups that often promote plant growth (e.g., *Burkholderia, Methylobacterium, Rhizobium*, and *Sphingomonas*; Table S8) [61–64]. In addition to richness and composition, the direct link between parasitism and duckweed population size (*r* = −0.76) indicated the role of other mechanisms. It is possible that phages reduced bacterial activity in the microbiomes via hijacking the molecular machinery of bacteria in the microbiomes to reproduce themselves [58]. It is also possible that like predation, parasitism by phages might drive the evolution of bacteria to develop defense traits, for example, toxin-antitoxin systems [65] that can potentially suppressing plant growth as an off-target effect [66]. Together, these results show predation and parasitism within microbiomes influence plant populations differently and highlight the need to elucidate the physiological and evolutionary mechanisms in understanding microbiome functions on plants.

Phosphorus is one of the most important limiting factors in freshwater ecosystems [67]. While both plants and microbes have been found to be important in mediating phosphorus cycling [68], our results disentangled their relative importance in phosphorus removal. Compared to microbiomes, duckweeds played a more important role in phosphorus removal in this system through direct phosphorus intake, as phosphorus is an important nutrient for plants. In addition to this mechanism, the photosynthesis of duckweeds might be able to change the environmental pH [69], which could influence phosphorus solubility and lower phosphorus availability [70]. While phosphorus is also important for bacterial growth as a cellular component for lipid and nucleic acid synthesis [71], bacterial abundance, richness, and composition did not influence phosphorus concentration directly (Fig. S2). Instead, the effects of bacterial richness and composition on phosphorus concentration were mediated through influencing duckweed population size. Furthermore, the opposing effects of predation and parasitism on phosphorus concentration were mediated by distinct mechanisms. While both predation and parasitism influenced phosphorus concentration via affecting duckweed population size, predation also reduced phosphorus concentration by altering other aspects of the microbiomes (the direct link, *r* = −0.31). This is likely because eukaryotic protists reduced phosphorus concentration by uptaking it through consuming bacteria, and this amount of phosphorus uptake is expected to be substantially higher than phages that transform a very small amount of bacterial phosphorus into own nucleotides [71]. Overall, our results emphasized that the opposing effects of predation and parasitism on phosphorus removal are primarily mediated via host plants, adding significantly to our knowledge of how different microbiomes (such as microbiomes from different origins, plant vs. environmental microbiomes, and natural vs. synthetic microbiomes) and within-microbiome trophic interactions mediate ecosystem functioning.

To summarize, our study provides the first experimental demonstration of how trophic interactions within microbiomes influence plant population size and ecosystem function. By leveraging the duckweed-microbiome symbiosis system, we show that not only do microbial predator-prey and host-parasite interactions operate in opposite directions in influencing plant population size and ecosystem function but also act via distinct mechanisms. It is important to distinguish between different types of trophic interactions when examining the role of trophic interactions for plant-microbiome symbiosis. These findings provide important insights into the host and ecosystem-level functions of microbiomes and call for the need of incorporating the knowledge of species interactions across multiple kingdoms and domains and the need of exploring diverse microbiome properties (including physiology and evolution) for better predicting the outcomes of complex host-microbiome systems.

## Data Availability

Raw experimental data will be deposited in Dryad. 16S rRNA amplicon sequencing data will be deposited in the NCBI Sequence Read Archive (SRA).

## Acknowledgments

This project is supported by Gordon and Betty Moore Foundation Symbiosis in Aquatic Systems Initiative (10635 to JT) and a research fund from Holden Arboretum (030869 to NW). MMT was supported by an NSF grant DEB (#1935410). We thank Jennifer Brum, Jae Kerstetter, and Yifeng Cao for assistance with the experiment, and Michael Hellberg and William Doerrler for allowing us using their lab space and equipment.

## Author Contributions

JT, NW, and MMT conceived the study. JT conducted the experiment and collected the data. NW conducted the data analyses and visualization. JT and NW wrote the manuscript. All authors contributed to revisions.

