## Supplemental figures for "Trophic interactions in microbiomes influence plant host population size and ecosystem function"

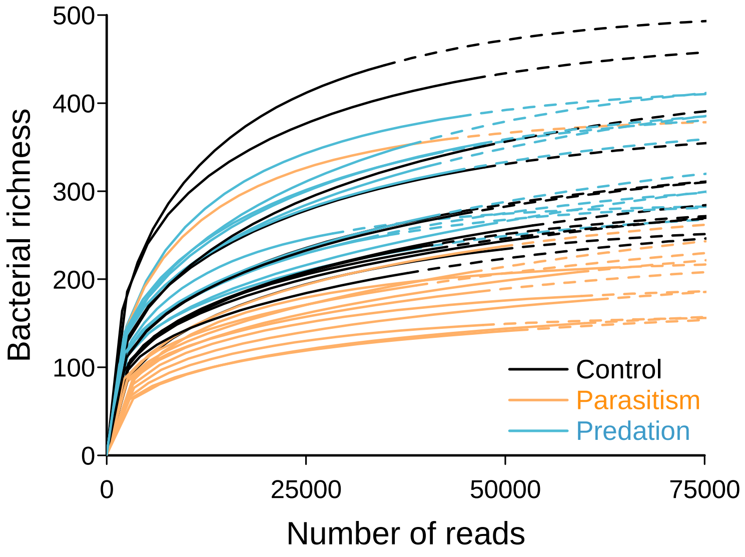


**Fig. S1 Rarefaction shows that the majority of bacterial richness was captured with our sequencing effort**. The number of sequencing reads (*x*-axis) obtained for bacterial microbiomes in individual experimental microcosms is represented by the solid portion of each colored line, whereas the dashed portion indicates extrapolation in the rarefaction analysis using the R package iNEXT.

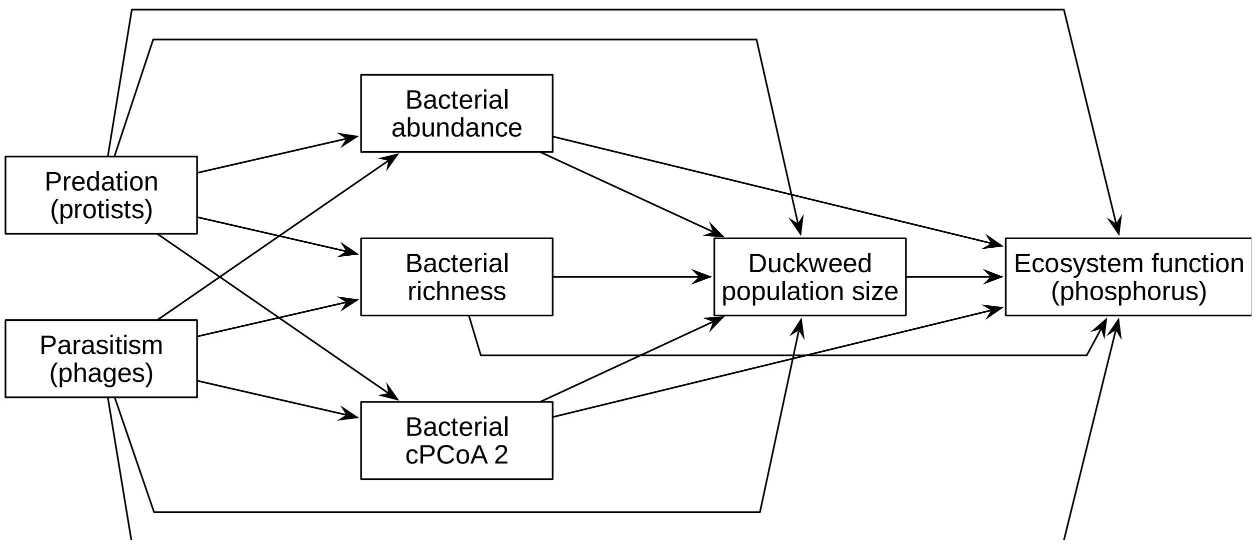


**Fig. S2 The full model of the structural equation modeling**
